# Quantification of *Aspergillus nidulans* Actin Dynamics during Early Growth and Septum Formation

**DOI:** 10.64898/2026.01.27.701996

**Authors:** Walker D Huso, Garrett Hill, Greeshma Tarimala, Jiyon Lee, Alexander G Doan, Junghun Lee, Kelsey Gray, Harley Edwards, Steven D Harris, Mark R Marten

## Abstract

Filamentous fungi have complex, three-dimensional growth patterns and a non-adherent nature, which can present challenges for live-cell imaging for quantitative assessment of dynamic cellular processes. To address these challenges, a live-cell imaging system has been modified to constrain the model fungus *Aspergillus nidulans* to growth in a single focal plane. This enables high-resolution time-lapse imaging of actin dynamics throughout development using a Lifeact actin marker. This system was used to perform kymographic analysis to quantify actin velocity and hyphal extension rates during early hyphal development. Results show two distinct growth phases: germ tube extension (0.58 μm/min) and hyphal extension (1.52 μm/min). Actin exhibited bi-directional transport along hyphae with biased movement toward the spore body. Actin was also observed re-localizing from hyphal tips to sites of septum formation indicating active redistribution of cytoskeletal resources based on cellular demands. This technological advancement overcomes longstanding limitations in fungal live-cell imaging and provides a new platform for quantitative systems-level analysis of mycelial development, offering new insights into the spatiotemporal coordination of cytoskeletal dynamics during filamentous growth.

## Introduction

The unique morphology of filamentous fungi is critical to their dual identity both as formidable pathogens and valuable commodity producers (*1*). The complex filamentous structure of these fungi requires intricate pathway regulation and sophisticated mechanisms to construct, maintain, and control their physiology. Actin, which plays an integral role in this process (*2*), is an abundant protein that has been well characterized throughout developmental phases across disparate fungi. It is instrumental in germination, septation, branching, hyphal extension, endocytosis, exocytosis, and vesicle trafficking (*3-7*). Globular actin (G-actin) organizes into higher order Filamentous actin (F-actin) structures including rings, patches, and cables all of which have unique roles in cellular development (*2*). Although well studied, actin dynamics have not been completely characterized or understood (*2*). Recently actin has been implicated in the survival response to cell wall stress (*8*) suggesting that actin-involved septal pathways play an important role in the survival mechanisms of *A. nidulans*. Specifically, the septation initiation network (SIN) is activated in response to cell wall stress and is hypothesized to recruit actin from sites of active growth in order to build more septa (*9*).

To study actin activity in microbial cells, fluorescent reporters have been added to truncated actin genes. This technique allows the fluorophore to localize with native actin proteins and the activity of actin to be tracked (*4, 6, 7, 10-12*). In filamentous fungi, this fluorescent tracking has revealed the role of actin in hyphal growth, septation, endocytosis as well as other activity and structures (*3, 4, 6, 10, 13-15*). These constructs have been used to identify critical cellular structures such as the contractile actin ring, cortical actin patch, and sub-apical actin web (*6*).

Many experimental approaches have been used to conduct live-cell imaging in filamentous fungi including: flow chambers, incubation chambers, agar block imaging, and complex fixation procedures(*2, 6, 10, 16, 17*). However, each of these techniques has significant limitations primarily involving the stages of growth which can be observed, and the length of time images can be captured. Filamentous fungi present particularly unique challenges for live cell imaging. They aggressively grow in 3 dimensions, are non-adherent, quickly outgrow microscope stages and have erratic growth characteristics making them difficult to precisely visualize and track over time (*17, 18*). This has resulted in significant limitations in existing imaging studies. Z-stack images have allowed for capture of physical details in 3-dimensionsal hyphae but require longer acquisition times incompatible with fluorescent imaging constraints. Alternatively, short acquisition times either limit imaging the focal plane to only capture specific features or a reduction in magnification to increase the focal depth. Until now constraining fungi in the z-dimension throughout developmental stages for long time-lapse sequences has not been possible. In this study, we adapted a commercially available flow chamber by using a novel gasket construct, to force fungal mycelia to be dimensionally constrained and grow in a single focal plane. This allows us to perform time course imaging of complete fungal entities with high time resolution. The Lifeact strain chosen for this experiment has an efficient *niiA* inducible promoter and expression of the Lifeact construct was shown to have negligible effect on growth rate at full induction(*6*). The novel flow chamber gasket assembly paired with the Lifeact-RFP reporter allows actin localization to be continuously tracked and measured throughout the growth of *Aspergillus nidulans*.

## Results and Discussion

The Lifeact strain used in this study is effective in allowing us to clearly visualize actin in different stages of development. For example, when this strain is grown in our flow chamber (Figure 1A-C), the subapical actin collar and subapical actin web (described by Shaw et. al. 2015) can be clearly observed(*11, 16, 19*). These same tip-localized structures were observed using confocal microscopy (Figure 1D), which also shows the cortical actin ring (Figure 1E) divided by a newly formed septum, forming a dual-actin ring. Growth in our flow chamber shows actin is clearly marked throughout fungal hyphae (Figure 1F) as actin patches and filamentous actin are generally visible where actin is traditionally expected to localize: actively growing tips, forming septa, and distributed along hyphae. Calcofluor white clearly marks the cell wall and mature septa after actin has re-localized.

**Figure 1.**
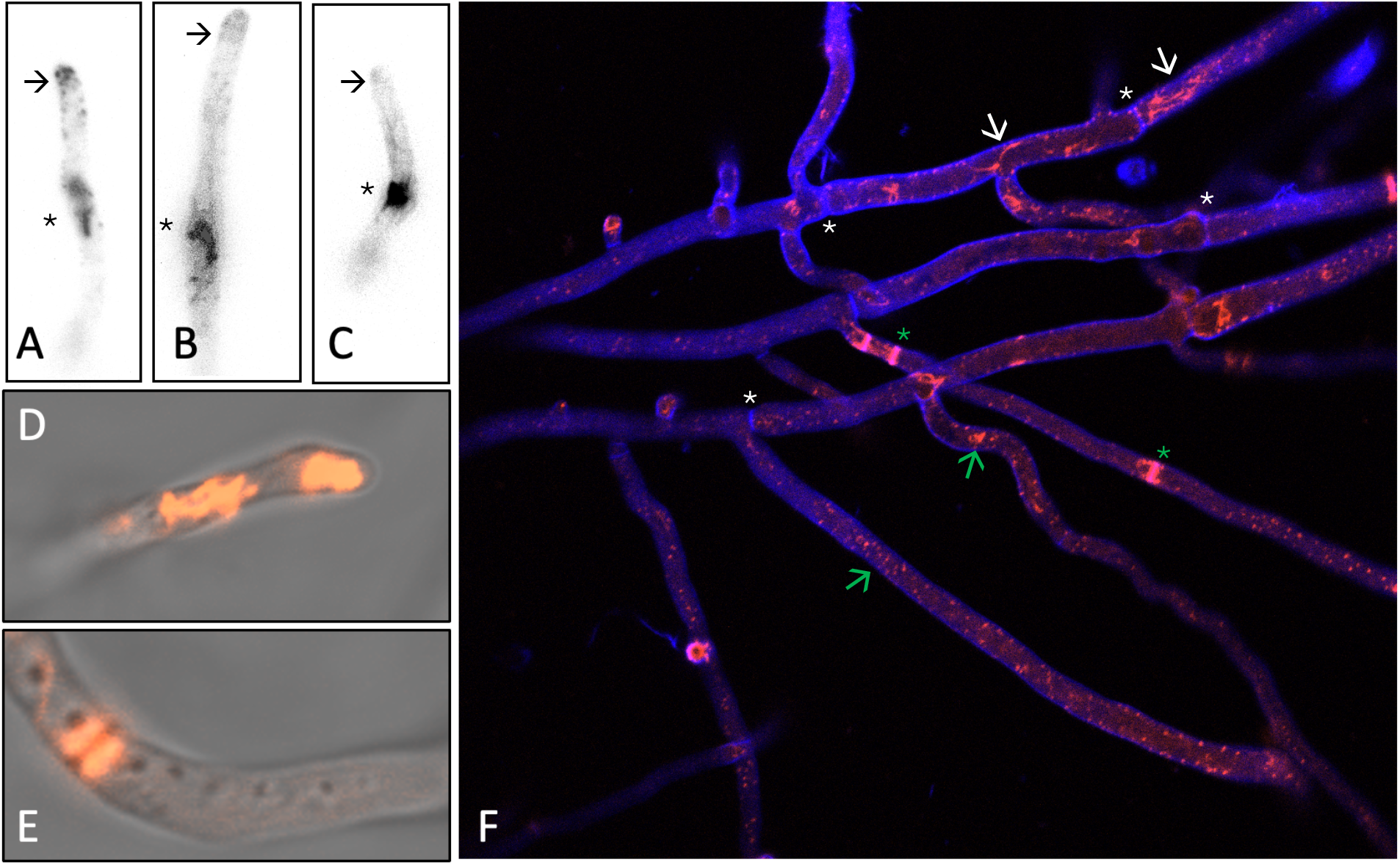
*Aspergillus nidulans* Lifeact-RFP construct can be used to highlight important developmental features. A-C) Actin can be seen localizing at the sub-apical collar (arrow) as well as the sub-apical actin web (star) D) Confocal microscopy shows actin localized at the hyphal tip of an actively growing hypha and E) a double contractile actin ring (CAR) divided by a newly developing septum. F) Confocal microscopy of the Lifeact strain stained with calcofluor white shows various stages of septation from actin aggregation and ring formation (green star) to mature fully formed septum (white star). Actin patches are seen distributed throughout (green arrows) as well as actin filaments (white arrows).

Figure 2 shows the Bioptechs flow chamber that was used to capture high resolution photos of actin movement during early hyphal development. To use the flow chamber, spores are adhered to the coverslip glass (Figure 2, A7) using Concanavalin A (Con A). The design of the flow chamber allows the Z-dimension of the growth chamber to be changed by varying the gasket thickness (Figure 2, A6). Through experiments with multiple different gasket materials of multiple thicknesses and using a number of different adhesives, we identified the combination described in the Materials and Methods section which allowed us to grow *A. nidulans* hyphae effectively in two dimensions (nominal 3 μm gap). As a result, the growing mycelium is always maintained in the same focal plane. Using this configuration we were able to measure relative fluorescence along hyphae to quantify changes in actin localization over time. In **Figure 2 G-I** the relative fluorescence is seen changing over time. Before septum formation (Figure 2G), fluorescence is mainly localized to the spore body and hyphal tip. During septum formation (Figure 2H) a spike in fluorescence intensity shows actin aggregates to the site of active septum development. Finally, after septum development is complete (Figure 2I) the actin dissipates and is once again seen localized to the spore body and hyphal tip.

**Figure 2.**
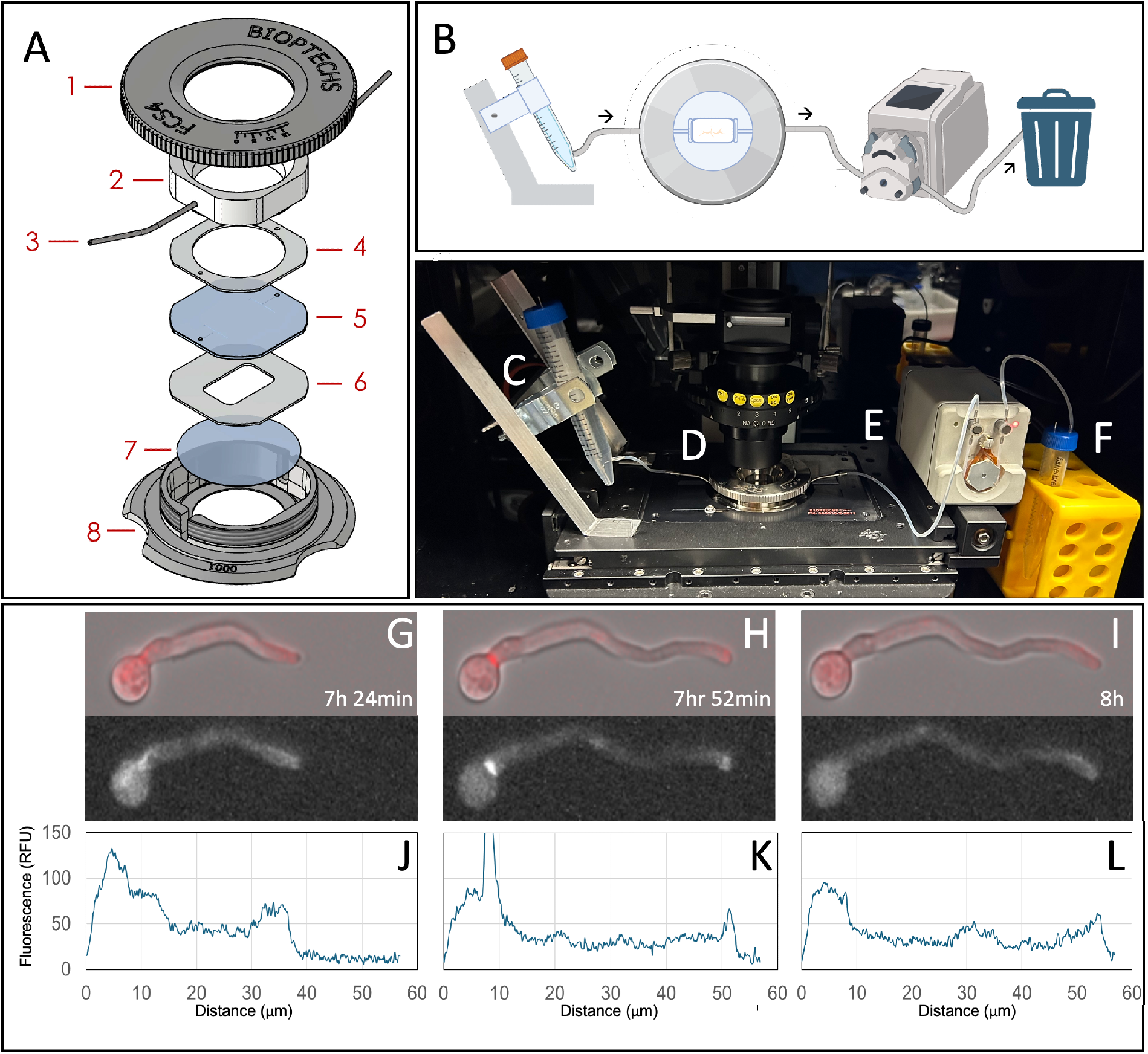
Bioptechs FCS4 flow chamber and experimental setup. (A) exploded diagram of the Bioptechs FCS4 Flow Chamber including: 1. chamber top 2. perfusion pressure plate 3. Perfusion inlet and outlet tubes 4. upper gasket 5. micro-aqueduct slide 6. lower gasket (cut from 3 μm mylar film) 7. 40 mm coverslip 8. self-aligning base and (not shown) an additional 0.05 mm silicon gasket between the coverslip and base to aid in sealing the mylar gasket. (B) Flow diagram showing the experimental setup of the flow chamber apparatus (arrows indicate media flow direction). (C) Experimental setup showing the designed media holder mounted to the microscope stage to minimize disturbance and tubing dead volume (D) flow chamber apparatus installed in the temperature controlled microscope enclosure (E) peristaltic pump with precise flow control and (F) waste container. Panels G-I are selected images of developing *A. nidulans* hyphae captured using flow chamber taken from timelapses of Lifeact strain (LQR3). Each pane includes merged (bright-field + fluorescent) and fluorescent images (i.e., Lifeact). (J-K) quantification of relative fluorescence along each hyphal element of images G-I respectively at indicated times showing the relative abundance of actin over the hyphal length.

To quantify actin dynamics and hyphal-tip extension rates over extended periods of early growth, time-course image sequences were captured and converted into kymographs(*20*) Representative kymographs (**Figures 3 and 4**) reveal the dynamic nature of actin behavior and demonstrate how actin re-localizes within the cell during growth. To create these figures, a line is drawn through the center of a specific hyphal element in a single image, and the average brightness along the length of the line is determined. This line is then represented by a single row of pixels in the kymograph. As a result, both time progression (i.e., top to bottom) and distance (i.e., left and right) are captured. Using these images, dynamic actin behavior can be quantified by measuring the angles of traces outlining Lifeact movement. Velocities (μm/min) are determined using trigonometry to calculate the slope. Actin movement is observed in both directions along the hyphae—from the spore body toward the tip and vice versa. Several distinct patterns of actin re-localization are observed. For example, in some cases actin moved from one hypha to another, across the spore body (Figure 3B, Velocity 6). In other cases, actin re-localized from the hyphal tip and sub-apical actin web (SAW) to sites of septal formation and actin ring assembly (Figure 4B, Velocity 3). Actin also returned to the hyphal tip following septum formation (Figure 4B, Velocity 4). These observations demonstrate the active role of actin in cellular processes and reveal how cells dynamically redistribute actin resources to support multiple developmental functions during normal growth.

**Figure 3.**
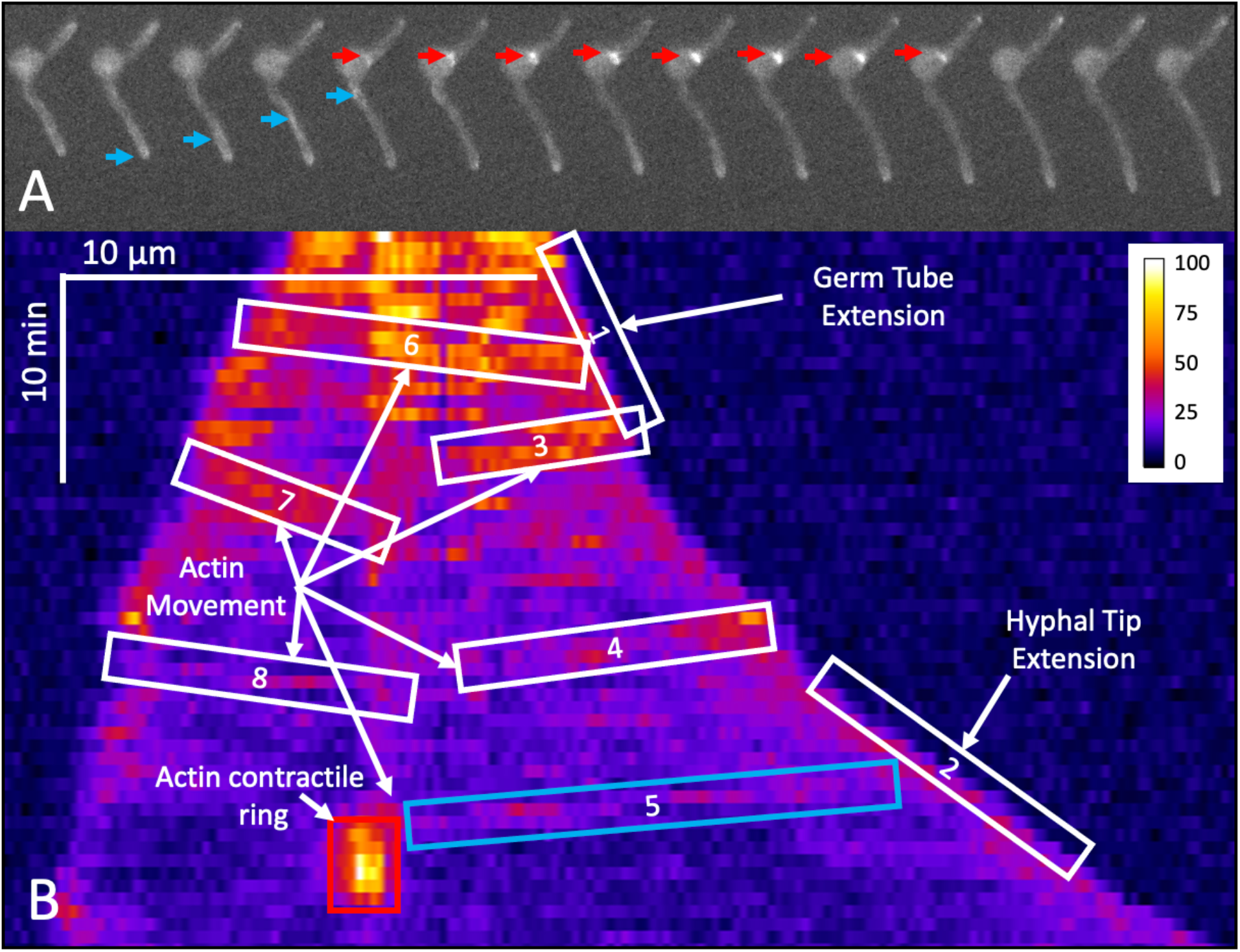
Representative kymograph showing early growth and development in *Aspergillus nidulans*. (A) Time-lapse images captured using the flow chamber show dynamic actin movement along hypha, toward spore body (blue arrows) and the formation and subsequent disappearance of the contractile actin ring (CAR) at the site of septum formation (red arrows) (B) Kymograph generated from 56-time lapse images (over 112 minutes) show actin localization during early-growth and development. Each row in the kymograph represents one timepoint. The growth of the tip is easily seen by the sloping front created on either side of the kymograph. The Inflection in the slope on the right side of the image represents an increase in the growth rate. The actin contractile ring is brightly marked by the Lifeact construct during septum formation (red box, corresponds to red arrows in A). Actin can be seen re-localizing from hyphal tip to the actin contractile ring (box 5, corresponds to blue arrow in A) prior to septum formation.

**Figure 4.**
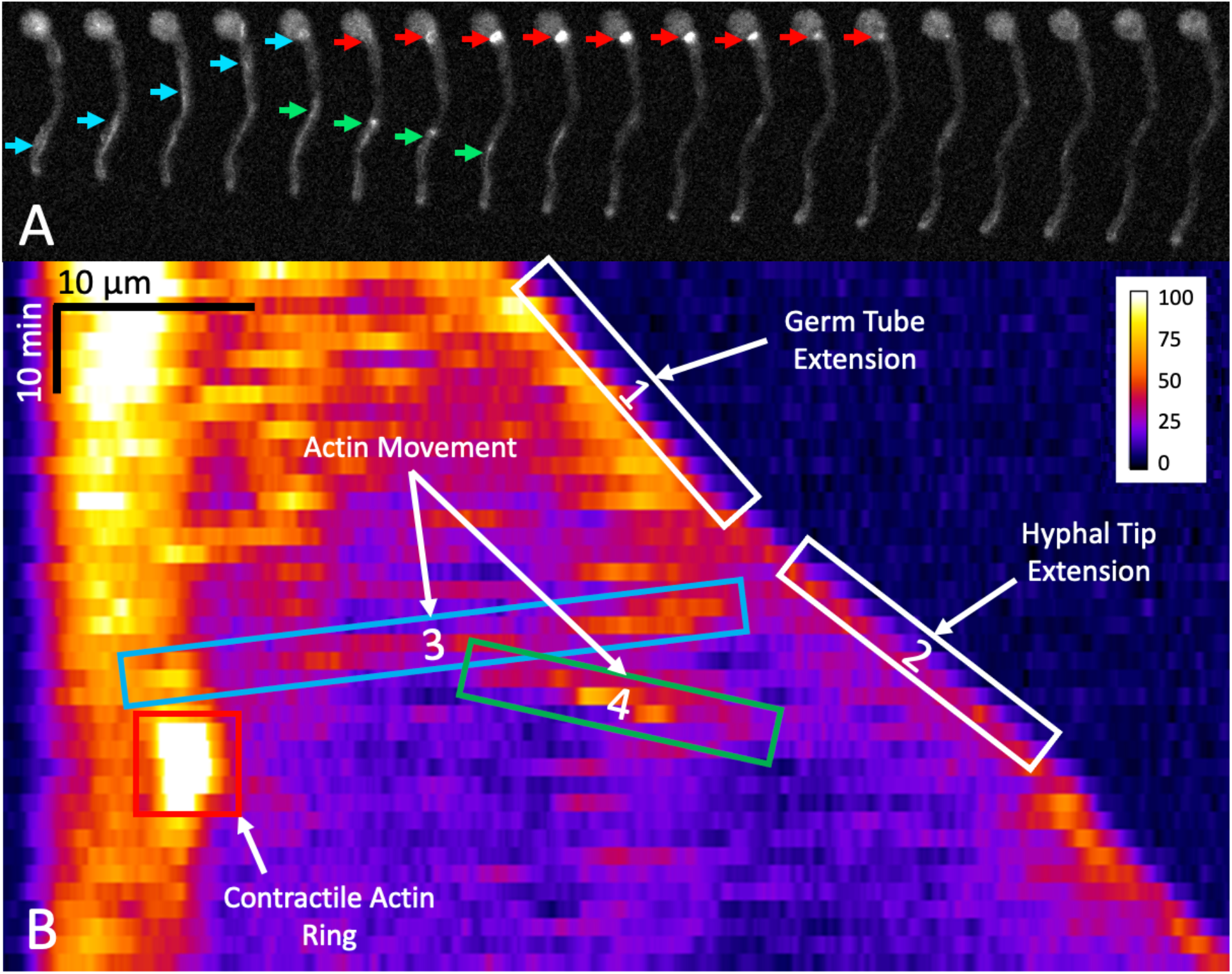
Kymographic analysis of early growth and development in *Aspergillus nidulans*. (A) Time-lapse images show actin re-localizing form the hyphal tip (blue arrows) towards the sight of septum development (red arrows). The formation of the CAR is indicated with red arrows. Green arrows show actin moving towards the hyphal tip. (B) Kymograph captures actin dynamics shown in (A) with matching color coding such that blue, green red boxes match similarly colored arrows. The hyphal growth rate is also captured by the slope.

The kymographs also capture changes in the tip extension rate. For example, immediately after germination hyphae grow relatively slowly, then transition to a higher growth rate (e.g., velocities 1 and 2 in Figures 3 and 4). This behavior was observed in all kymographs as a unique inflection point. This implies there is a change in growth regimen from germ tube extension to hyphal growth, resulting in two distinct growth regimens in early fungal development, which is consistent with literature(*21*). These two growth phases have been described as the germ tube extension rate and the hyphal extension rate. During the initial stages of development and prior to first septal formation, hyphal growth was observed to proceed at a significantly slower rate. In the kymographic analysis (Figure 3, 4), a sharp inflection point was observed in the tip velocity shortly preceding septation indicating a rapid acceleration in the tip extension rate. The difference in germ tube extension and hyphal extension was even more pronounced after the analysis shown in Figure 5 and statistical testing. The germ tube extension rate averaged 0.58 μm/min, while the hyphal extension rate averaged 1.52 μm/min, nearly 3 times faster. These growth rates were substantially higher than the reported growth rate in previous studies using the same Lifeact strain [6], suggesting that this flow chamber system provides more favorable growth conditions compared to those experienced during conventional agar-block imaging.

**Figure 5.**
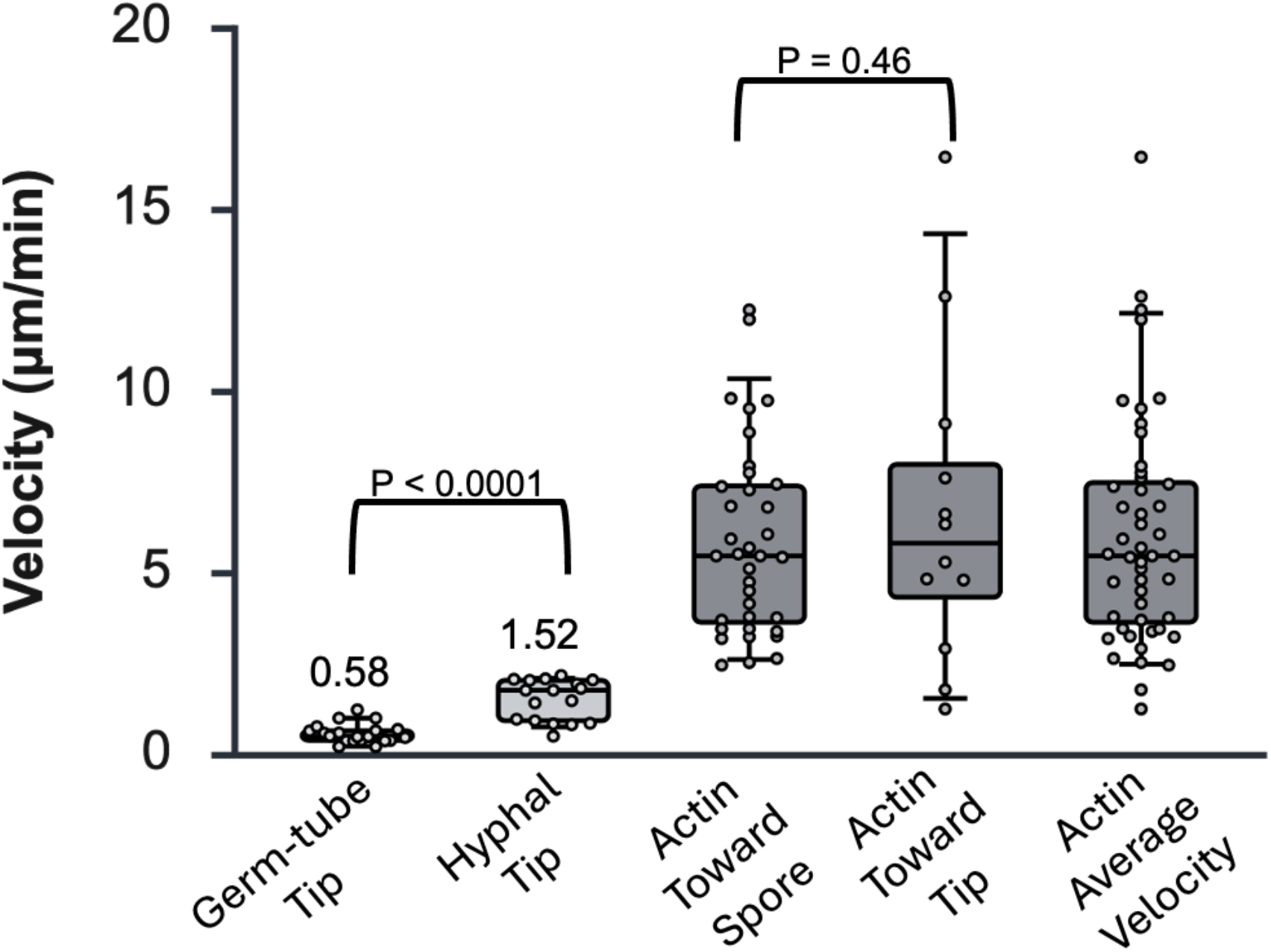
*A. nidulans* tip growth-rate and actin velocity during early growth and development. The germ tube extension rate is shown to be significantly less than the hyphal extension rate. Actin moves bidirectionally (to and from the spore body) with no significant difference in average velocity. However, actin movement toward the spore body was found to be much more frequent than movement away from the spore body.

Actin movement was observed continuously throughout growth and development, with bidirectional transport along hyphal elements. In several instances, actin was observed re-localizing from actively growing hyphal tips to sites of septal development, where it apparently contributed to actin contractile ring formation. The velocity of actin movement was consistently faster than tip growth rates. Although both anterograde and retrograde movement of actin relative to the growing tip proceeded at a statistically similar rate, actin transport away from the tip was much more common. This indicates that proportionally more actin is produced near the tip before being transported to areas of active growth. This coincides with mRNA localization studies in other filamentous fungi that shows actin mRNA’s localize to the apical region (*13, 22*). Interestingly, actin was also documented moving through the lumen of the spore body between different hyphal elements. These observations imply that actin is dynamically re-purposed and re-allocated based on cellular demands and recruited to sites of active development in response to stimuli and cellular requirements.

Altogether, this advancement in flow chamber technology opens significant new possibilities for understanding fungal development and transitions between developmental phases. Highly detailed visualization of long-term growth will be valuable in understanding how developmental processes are connected and how fungi grow and mature. The combination of available genetic fluorescent markers and the genetic tractability of filamentous fungi, particularly *Aspergillus nidulans*, makes the future applications of this technology exceptionally broad. This approach introduces a powerful new tool for comprehensive studies of mycelial development and growth dynamics.

## Materials and Methods

### Strains, Media, and Growth Conditions

In this study, the model fungus, *Aspergillus nidulans*, strain LQR3 was used (pyrG89::niiA(p)::Lifeact::TagRFP; pyroA4; wA3; veA1) (*6*). Frozen stocks were spread on MAGV plates (2% malt extract, 1.5% agar, 2% glucose, 2% peptone, and 1 ml/L Hutner’s trace elements and vitamin solution) and incubated for 3 days at 28 °C (*23*). Spore lawns were harvested into 10ml sterile diH20 and diluted to a concentration of 1×10^5 spores/ml and used to inoculate the flow chamber. Spores were adhered to the cover glass using concanavalin A solution (20 mL of PBS, 100 μL 132 mM CaCl2, and 10.7 mg Concanavalin A) (*24*). For imaging experiments, nitrogen deficient minimal media (MM-) was amended with 10mM KNO_3_ to ensure full induction of the *niiA* promoter (*6*).

### Flow Chamber

The Bioptechs FCS4 (Bioptechs Inc., Butler, PA) flow chamber was used for live-cell imaging. Each component was autoclaved separately before assembly to prevent contamination. To assemble the system, a drop of Concanavalin A solution was placed on to 40mm Borosilicate coverslips. A 3µm mylar film (Premier Lab Supply, Port St. Lucie, FL) was then placed on top and smoothed onto the coverslip forcing out excess ConA solution. The excess mylar film was then trimmed around the edge and the coverslip dried. The internal growth chamber geometry was designed using CorelDraw software. The 1×2cm design was cut from the center of the mylar coating the coverslip using a CO2 laser cutter (630−680 nm, max output is 5 mW, Class 3R, 2.0 lens module, Universal laser systems, Arizona) exposing the ConcA treated coverslip. Spore solution was pipetted onto the coverslip and dried allowing spores to adhere to the ConcA. The flow chamber was assembled, connected to an upstream media reservoir and a downstream waste receptacle. media flow was controlled using an P720 peristaltic pump (Instech, Plymouth Meeting, PA) and 0.015” ID silicon tubing (Instech BSILT015). The Flow rate was set to ∼17µl/min.

### Imaging

An Olympus IX-81 inverted fluorescence microscope (Olympus, Tokyo, Japan) equipped with a fluorescent Lumencor SOLA Light Engine and Hamamatsu ORCA-spark camera was used to capture images. 40x magnification was used to capture timelapse images. 100x oil immersion was used to capture highly detailed static images. After the flow chamber was assembled and filled with the full-induction media, it was pre-incubated for 6hrs at 28°C. After 6hrs Cell-Sens software was used to take time-lapse photos every 2 min. Bright-field images were taken with exposures of 10-50µs. A Texas Red filter was used to capture the Lifeact fluorescence. These images were taken with exposure of 5s. Confocal images were captured on a ZEISS LSM 900 Confocal with Airyscan 2 microscope with a 63x oil immersion lens using agar block imaging techniques(*6, 25*).

### Image analysis

FIJI image analysis software (https://imagej.net/software/fiji/) (*26*) was used to analyze time-lapse image sequences. The Kymograph builder plugin was used to generate kymographs based on region of interest (ROI) drawn along the length of the fungal element. The width was set to 19 pixels to match the average width of the hyphae. The angle of the slopes of interest (actin movement and hyphal tip) in the kymographs were measured using the Angle Tool. The slope was then calculated using arctan(θ) and converted from pixel/pixel to µm/min. The unit conversions are 0.1465µm/px on the x-axis and 2min/px for the y-axis.

## Acknowledgements

This work was supported by the National Science Foundation (Awards 2006189, 2527369, 2527370). We gratefully acknowledge Dr. Tagide deCarvalho and the Keith R. Porter Imaging Facility for the use of the Confocal microscope. We also acknowledge Dr. Govind Rao (UMBC, Center for Advanced Sensor Technology) for use of the laser cutter for preparing the mylar coverslip seals.

## Notes

### Competing Interest Statement

The authors have declared no competing interest.

